# Single-shot Mesoporous Silica Rods Scaffold for Induction of Humoral Responses Against Small Antigens

**DOI:** 10.1101/2020.03.17.993808

**Authors:** Maxence O Dellacherie, Aileen Li, Beverly Y Lu, Catia S Verbeke, Luo Gu, Alexander G. Stafford, Edward J. Doherty, David J. Mooney

## Abstract

Vaccines have shown significant promise in eliciting protective and therapeutic responses. However, most effective vaccines require several booster shots, and it is challenging to generate potent responses against small molecules and synthetic peptide antigens often used to increase target specificity and improve vaccine stability. As continuous antigen uptake and processing by APCs and persistent toll-like receptor (TLR) priming have been shown to amplify antigen specific humoral immunity, we explored whether a single injection of a mesoporous silica micro-rod (MSR) vaccine containing synthetic molecules and peptides can effectively generate potent and durable antigen-specific humoral immunity. A single injection of the MSR vaccine against a gonadotropin-releasing hormone (GnRH) decapeptide elicited highly potent anti-GnRH response that lasted for over 12 months. The MSR vaccine generated higher titers than bolus or alhydrogel alum vaccine formulations. Moreover, a MSR vaccine directed against a Her2/neu peptide within the Trastuzumab binding domain showed immunoreactivity to native Her2 protein on tumor cell surface and, when directed against nicotine, generated long-term anti-nicotine antibodies. Mechanistically, we found that the MSR vaccine induced persistent germinal center (GC) B-cell activity for more than 3 weeks after a single injection, generation of memory B cells, and that at least 7 days of immunostimulation by the vaccine was required to generate an effective humoral response. Together, these data suggest that the MSR vaccine represents a promising technology for synthetic antigen vaccines to bypass the need for multiple immunizations and enhance long-term production of antibodies against endogenous antigens in the context of reproductive biology, cancer, and chronic addiction.

## 1. Introduction

Immunotherapies are rapidly becoming a standard approach for cancer and chronic infectious disease treatment ^[1]^. Vaccines can induce strong protective and therapeutic responses against many forms of live and attenuated virus, toxoids, irradiated autologous tumor cells and protein subunits. However, most effective vaccines require several boosts after the primary immunization ^[2]^. These dosing regimens are costly, increase hospital/clinic visitation for patients and are especially challenging in regions where limited healthcare access poses a major logistical barrier to disease treatment and management ^[2,3]^. An effective single-shot vaccine could overcome these limitations. Most approaches for single-shot vaccines in clinical testing to date have used depots of antigens that were released in a controlled manner for months after immunization ^[2,4–6]^. However, passive diffusion of the antigen and adjuvant to the lymph nodes alone is likely unable to maximally and rapidly stimulate immune cells and provide life-long benefit. Indeed, besides B-cell stimulation by antigen, humoral responses development requires a complex immune cascade involving antigen-presenting cells (APC) activation, subsequent T-helper cell generation and finally cognate T-cell/B-cell interactions. However, these steps that occur in parallel during infection are not easily recapitulated by current vaccine technologies. Therefore, achieving a robust and lasting immune response after a single injection of a vaccine remains an important challenge.

Small molecules and peptides have many potential advantages as antigens but generating potent responses is challenging. The use of small antigens in vaccine can help focus the immune response with high epitope specificity and biological activity while avoiding off-target effects. For instance, a synthetic peptide vaccine against subtypes of influenza using Keyhole limpet hemocyanin (KLH) as a carrier protein generated anti-hemagglutinin antibodies and neutralizing activity ^[7]^. Peptides can be particularly advantageous for vaccine development as they can be chemically synthesized quickly and with low variability. Their production can be easily up-scaled and they have good safety and stability ^[8]^. In contrast to the time and cost-intensive approaches that require the production of recombinant protein, attenuated virus or cancer cell culture, synthetic peptides can expedite the manufacturing of personalized cancer vaccines and ensure the timely development of vaccines against new pandemics ^[9–11]^. Peptides are also readily presented by antigen presenting cells (APCs) to induce subsequent effector T cell and humoral responses. However, synthetic peptide vaccines generally cannot confer long-lived, therapeutic benefit. This limitation is likely caused by a combination of 1) short peptide half-life due to rapid systemic clearance, 2) weak immunogenicity due to lack of a co-stimulatory signal and either absent or suboptimal T-helper epitopes co-presentation, and 3) absence of B-cell receptor (BCR) cross-linking due to lack of multivalency^[8,12,13]^.

An example of an attractive target for vaccination is gonadotropin-releasing hormone (GnRH), a decapeptide hypothalamic hormone that controls male spermatogenesis, and female estrogen and follicular development ^[14]^. GnRH determines both fertility and sexual behavior and is highly conserved among mammals. GnRH activity has also been implicated in the progression of several types of malignancies including prostate, breast and ovarian cancers ^[15–17]^. Blocking GnRH activity using androgen deprivation therapies (ADTs) and gene therapy have shown pre-clinical promise, but such therapies have strong toxic side effects and lack long term potency ^[12,16]^. Active immunizations against GnRH using various carrier proteins such as KLH, ovalbumin (OVA) and diphtheria toxoid (DT) have been shown to decrease sexual organ function, reduce testosterone levels in patients with advanced prostate cancer and reduce reproductive capability in feral animals as a humane alternative to surgical desexing ^[16–18,20]^. However, these vaccines require multiple immunizations and the effects are variable and not potent enough to produce long-term immune-castration ^[21]^.

Here we propose a single injection vaccine platform using a mesoporous silica micro-rod (MSR) vaccine system to generate high antibody titers against synthetic peptide and other small antigens. Biomaterials have shown substantial potential to integrate with conventional vaccine approaches by, for example, extending the half-life of immune-modulating drugs and controlling the presentation of adjuvants and antigens ^[22–24]^. In particular, biomaterial scaffold vaccines have been demonstrated to locally recruit and program host antigen presenting cells (APCs) to induce effective innate and adaptive immunity ^[25,26]^. Using this approach, an implantable poly(lactide-co-glycolide) (PLGA) based scaffold cancer vaccine was shown to induce complete melanoma regression in subsets of mice ^[27]^. To eliminate the need for surgical implantation, we recently developed a new approach of injectable scaffolds using mesoporous silica rods (MSR) microparticles ^[28,29]^. We demonstrated that MSR microparticles could spontaneously assemble into a 3D scaffold after subcutaneous injection; the macropores formed by random particle stacking allowed for cell infiltration and active cell-scaffold interactions. The controlled release of GM-CSF and CpG from the MSR vaccine modulated host APC infiltration into the vaccine and, consequently, continuously programmed host APCs. Continuous antigen uptake and processing by APCs and persistent toll-like receptor (TLR) priming have been shown to amplify antigen specific humoral immunity. Overall, we reasoned that concomitant release of antigen from MSR scaffold and local APC activation could enable continuous B-cell activation in the lymph node and enhance cognate T-helper cell priming (Figure 1). Therefore, we hypothesized that a single injection of the MSR vaccine could generate robust and long-lasting humoral immunity and enhance the response compared to traditional vaccine approaches. We focus on GnRH as a model antigen, but also explore this concept with a small molecule epitope nicotine and a Her2 peptide epitope within the Trastuzumab-binding domain.

**Figure 1.**
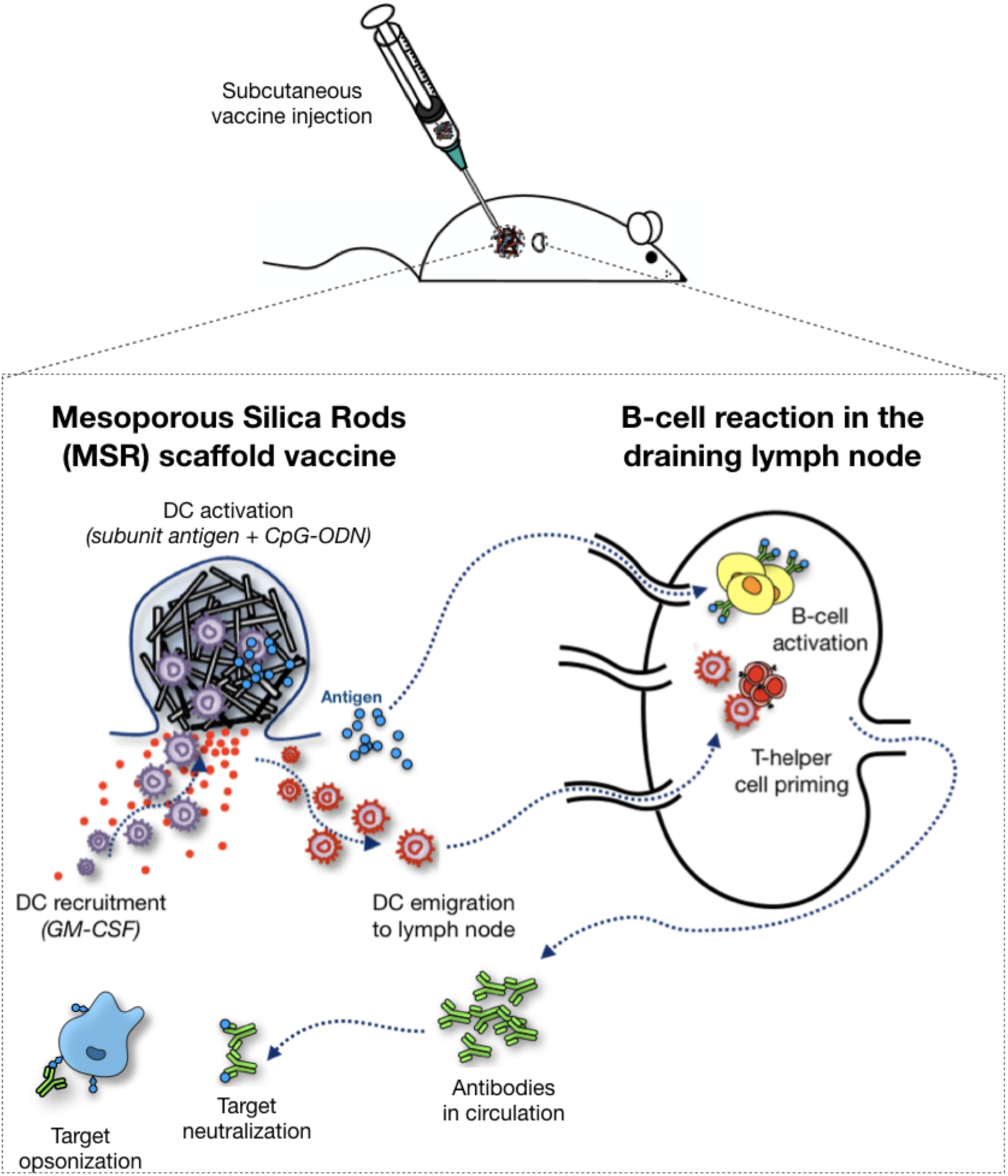
Mesoporous Silica Rods vaccines for humoral response generation. Mesoporous Silica Rods loaded with GM-CSF, CpG-ODN and subunit antigens are injected subcutaneously to form a 3D immunomodulating scaffold to produce antibodies against small antigen targets. MSR scaffold serves two purposes (1) as a long-term depot for sustained antigen release and presentation to B-cells in the draining lymph node. (2) as a local niche for dendritic cell (DC) recruitment where they pick up antigen and get activated by TLR-9 agonist CpG. Mature DCs can migrate to the lymph node where they prime antigen-specific T-cells that support the development of the humoral response.

## 2. Results

### 2.1 Single injection of MSR vaccine with GnRH peptide conjugated to OVA (GnRH-OVA) induced higher and more durable anti-GnRH humoral response compared to bolus vaccines

To evaluate whether a CD4 helper T cell epitope was necessary to generate an anti-GnRH response, we incorporated either free GnRH peptide or GnRH peptide conjugated to the carrier protein ovalbumin (OVA) into the MSR vaccine. MSR vaccines were formulated to contain 1µg GM-CSF and 100µg CpG-ODN, as previously described, and either 100µg free GnRH or 100µg GnRH conjugated to 200µg OVA (GnRH-OVA) (Figure 2a). GnRH-OVA conjugation was confirmed using mass spectrometry (Figure S1). The loading efficiencies of free GnRH and GnRH-OVA onto the MSRs were 91 ± 2% and 85 ± 1%, respectively (Figure S2). Vaccines were injected into the subcutaneous tissue of mice, and antibody titers were subsequently measured using ELISA. Strikingly, a single immunization with the MSR vaccine containing GnRH-OVA elicited strong titers of anti-GnRH IgG1 (Figure 2b) and IgG2a (Figure 2c) serum antibody for over 12 months. In comparison, immunization with MSRs loaded with unconjugated GnRH did not elicit detectable anti-GnRH titer. To confirm the serum anti-GnRH antibody recognized the GnRH peptide sequence specifically, serum from animals vaccinated with the MSR GnRH-OVA vaccine were tested using an ELISA coated with GnRH peptide or a scrambled peptide (CRSYGPLHEWG) (Figure S3a). The serum antibody recognized the GnRH peptide in a dilution-dependent manner but did not recognize the scrambled peptide. Finally, to confirm that the anti-GnRH antibody response was specific to the MSR GnRH-OVA vaccine, we immunized mice with MSR vaccines containing 200µg OVA (MSR OVA). Serum from animals vaccinated with the MSR OVA vaccine did not elicit any detectable anti-GnRH IgG1 (Figure S3b) or IgG2a (Figure S3c) titers.

**Figure 2:**
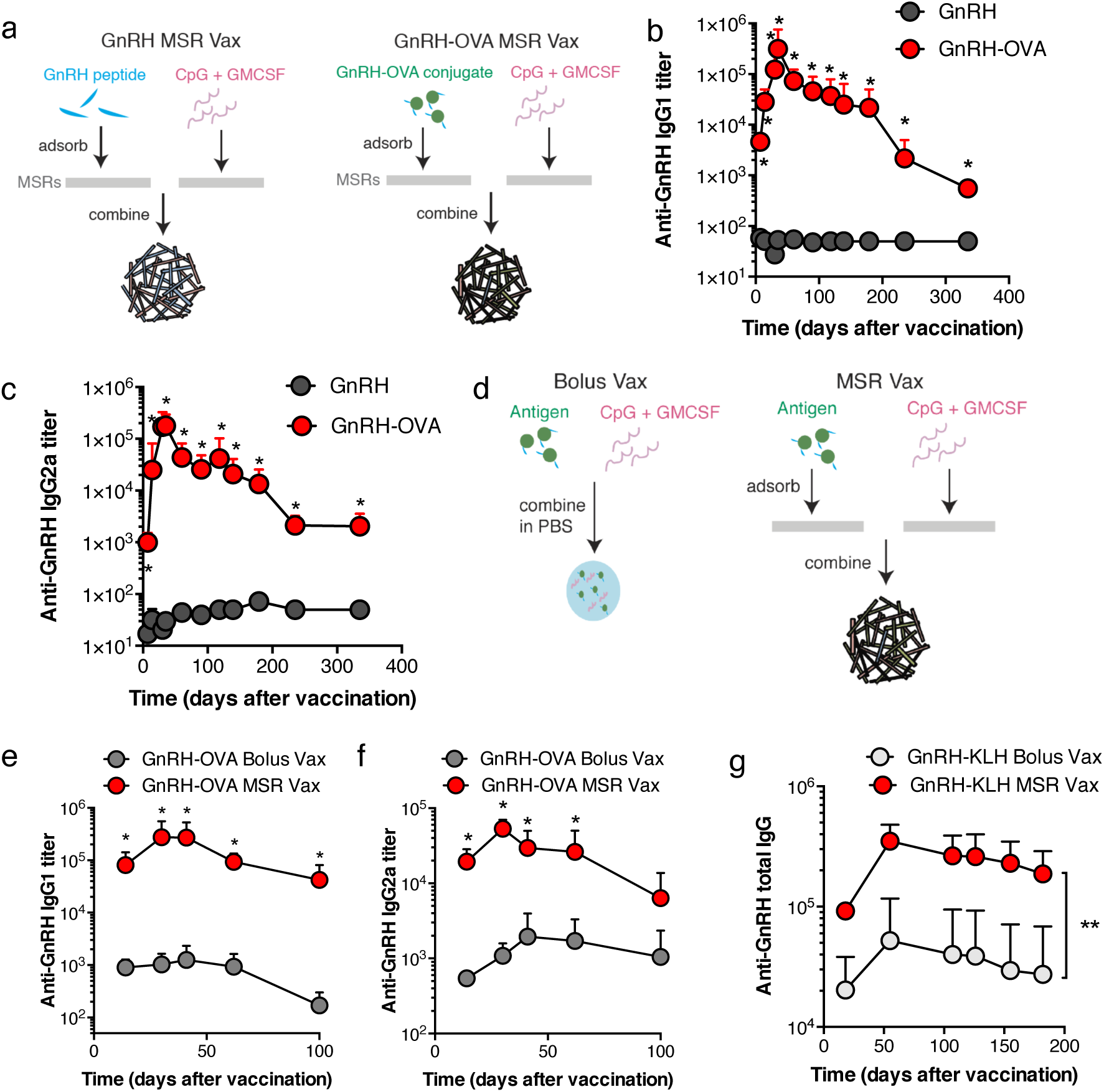
Single injection of MSR vaccine with GnRH-OVA induces durable anti-GnRH peptide serum antibody response, and higher titer than bolus formulations. (a) Schematic of MSR vaccines formulations using free GnRH peptide (left) or GnRH-OVA (right). (B-C) ELISA analysis of sera GnRH-specific IgG1 (b) or IgG2a (c) after immunization with MSR vaccines containing free GnRH or GnRH-OVA (mean and s.d., n = 4) * indicates p < 0.05. (d) Schematic of Bolus vaccine (left) and the MSR vaccine (right). (E-F) ELISA analysis of sera GnRH-specific IgG1 (e) or IgG2a (f) after immunization with MSR vaccines loaded with GnRH-OVA (GnRH-OVA MSR Vax) or bolus vaccine (GnRH-OVA Bolus Vax). (mean and s.d., n = 4) * indicates p < 0.05. (g) ELISA analysis of sera GnRH-specific total IgG after immunization with MSR vaccine loaded with GnRH-KLH (MSR Vax) or bolus vaccine (Bolus vax)). mean and s.d., n = 4 ** indicates p < 0.01

We next investigated whether the MSR vaccine can elicit higher antibody titers compared to traditional bolus immunizations (Figure 2d). Mice were again immunized with a single injection of the MSR vaccine loaded with 1µg GM-CSF, 100µg CpG-ODN and 100µg GnRH conjugated to 200µg OVA or a single injection of a bolus vaccine containing 1µg GM-CSF, 100µg CpG-ODN and 100µg GnRH conjugated to 200µg OVA in PBS (GnRH-OVA Bolus Vax) (Figure 2d). The MSR vaccine induced significantly higher anti-GnRH IgG1 (Figure 2e) and IgG2a (Figure 2f) antibody titers compared to the bolus vaccine. The effect was also durable, lasting for at least 100 days post immunization. Impressively, the MSR vaccine induced rapid onset of antibody production. On day 14, the titer elicited by the MSR vaccine was 35 fold and 80 fold higher than that elicited by the bolus vaccine for IgG2a and IgG1 subtypes, respectively. Keyhole Limpet Hemocyanin (KLH) is the most commonly used carrier protein immunogen to induce antibody production, as it is highly immunogenic due to its large molecular weight and repetitive structure, which allow for high valency antigen conjugates ^[30]^. Using GnRH-KLH conjugate as the antigen, MSR vaccine again significantly enhanced the total serum anti-GnRH IgG response compared to a bolus KLH-GnRH vaccine (Figure 2g). At the peak of the response, the titer elicited by the MSR vaccine was nearly 10 fold higher than that elicited by the bolus vaccine.

### 2.2 Single injection of MSR vaccine induced persistent Germinal Center (GC) activity

A strong humoral response depends on activated B cells forming GCs in secondary lymphoid organs, where they further undergo somatic hypermutation and isotype switching, and eventually differentiate into plasma cells that can rapidly produce antibodies upon secondary encounter with the antigen ^[31]^. Therefore, we investigated whether a single injection of the MSR GnRH-OVA vaccine can stimulate enhanced GC formation compared to the conventional bolus vaccination. The MSR GnRH-OVA vaccine induced a prolonged immune response, as indicated by the increased total cell number in the dLN from 7 days to 25 days after immunization (Figure 3a). In contrast, although the bolus GnRH-OVA vaccine enhanced dLN cell number on day 7, the response rapidly dissipated. Next, the percentage and total cell number of GC B cells in the dLN were quantified using the markers B220^+^GL7^+^ (Figure 3b and 3c) and B220^+^GL7^+^PNA^+^ (Figure S4a and S4b, primary gating plots shown in Figure S5) using flow cytometry. By day 7 after immunization, both the MSR vaccine and the bolus vaccine led to an enhanced GC response. Notably, the enhanced GC response in the MSR vaccinated animals persisted until after day 25. The number of GC B cells in the MSR vaccinated mice was approximately 10 fold higher than those in the bolus vaccine treated mice on days 14 and 25. In contrast, the GC response in the bolus vaccinated animals sharply dissipated after 7 days. These findings were further supported by immunofluorescence staining of GC B cells in the dLNs over time (Figure 3d).

**Figure 3:**
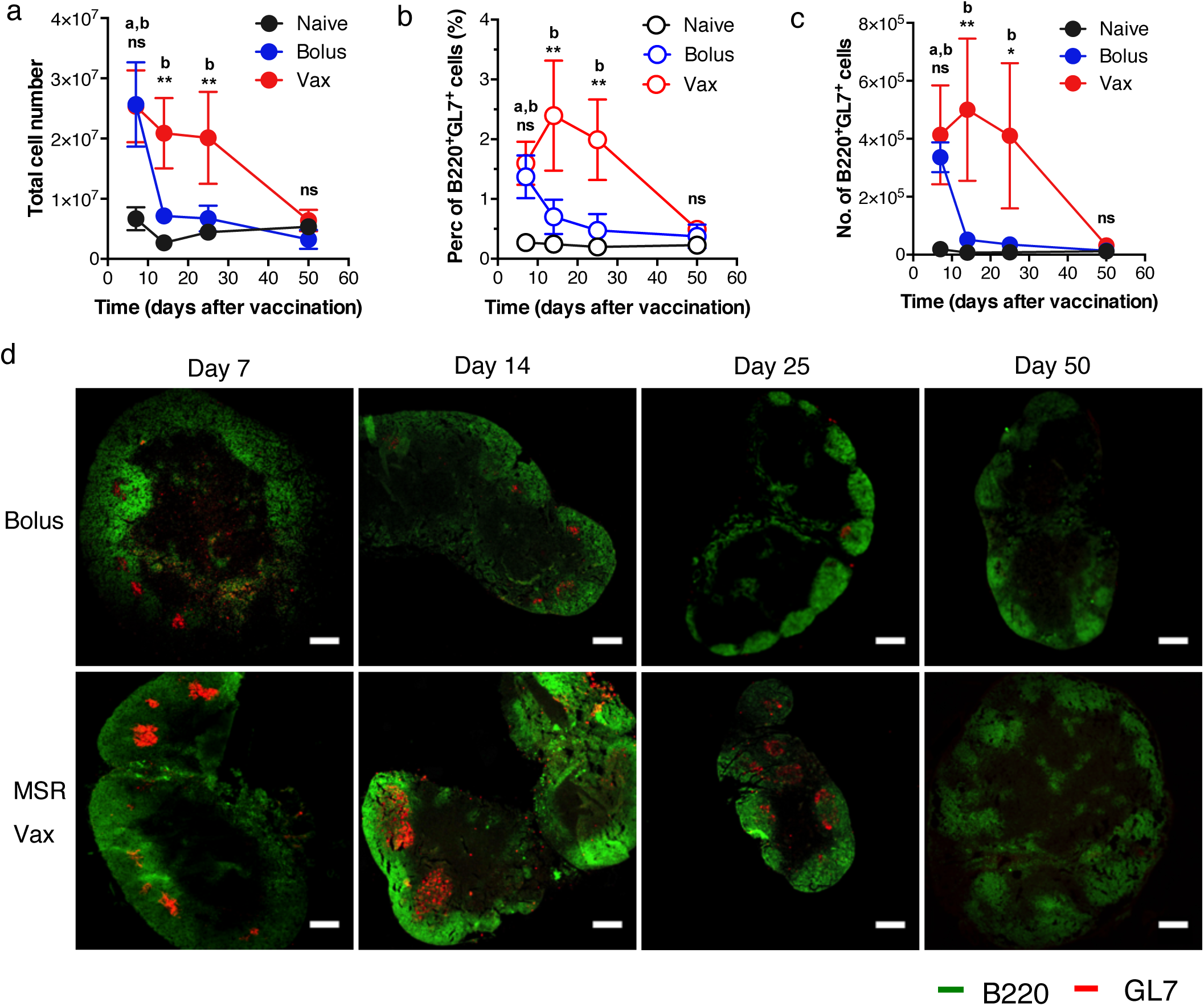
MSR vaccine with GnRH-OVA induces persistent germinal center formation. (a) Total number of cells in the dLN of untreated mice (Naïve) or after immunizing with the MSR vaccine (vax) and bolus formulations of the vaccine (Bolus). (b) Percentage and (c) number of B220^+^GL7^+^ GC B cells in the dLN (c) Histological staining of dLN sections stained with anti-B220 and anti-GL7. scale bar = 300µm.. (mean and s.d., n=4) a indicates p < 0.05 between Naïve and Bolus group, b indicates p<0.05 between Naïve and Vax, * indicates p <0.05, ** p<0.01 and ns, no significance between Bolus and Vax

### 2.3 Single injection of MSR vaccine improves memory B-cell generation

Efficient prophylactic vaccination requires the induction of long-term immunological memory to protect against pathogen encounter at a later time. Memory B-cells generated by vaccination can persist for years and be reactivated upon antigen encounter to produce rapid antibody responses ^[31]^. Therefore, we compared the capacity of bolus and MSR-based vaccines to induce such responses. To detect the rare memory B-cells generated by the vaccine, we used fluorescent protein phycoerythrin (PE) as both the antigen and the detection reagent (Figure 4a) ^[32]^. 10-12 weeks after immunization, several weeks after all vaccine components had cleared (Figure 4b), cells from the draining lymph node (dLN), spleen and bone marrow were stained with PE, enriched for PE-binding cells using anti-PE magnetic and finally, PE-specific B-cells were detected by flow cytometry (Figure S6). Compared to naïve mice, MSR-immunized mice showed greatly increased numbers of PE^+^B220^+^ B-cells in the draining lymph node and spleen. By contrast, bolus immunization did not provide a significant boost in antigen-specific B-cells compared to the naïve baseline. No differences were observed in the bone marrow for any of the groups (Figure 4c). Overall, this data illustrates that MSR vaccine can enhance memory B-cell generation compared to traditional bolus vaccination.

**Figure 4:**
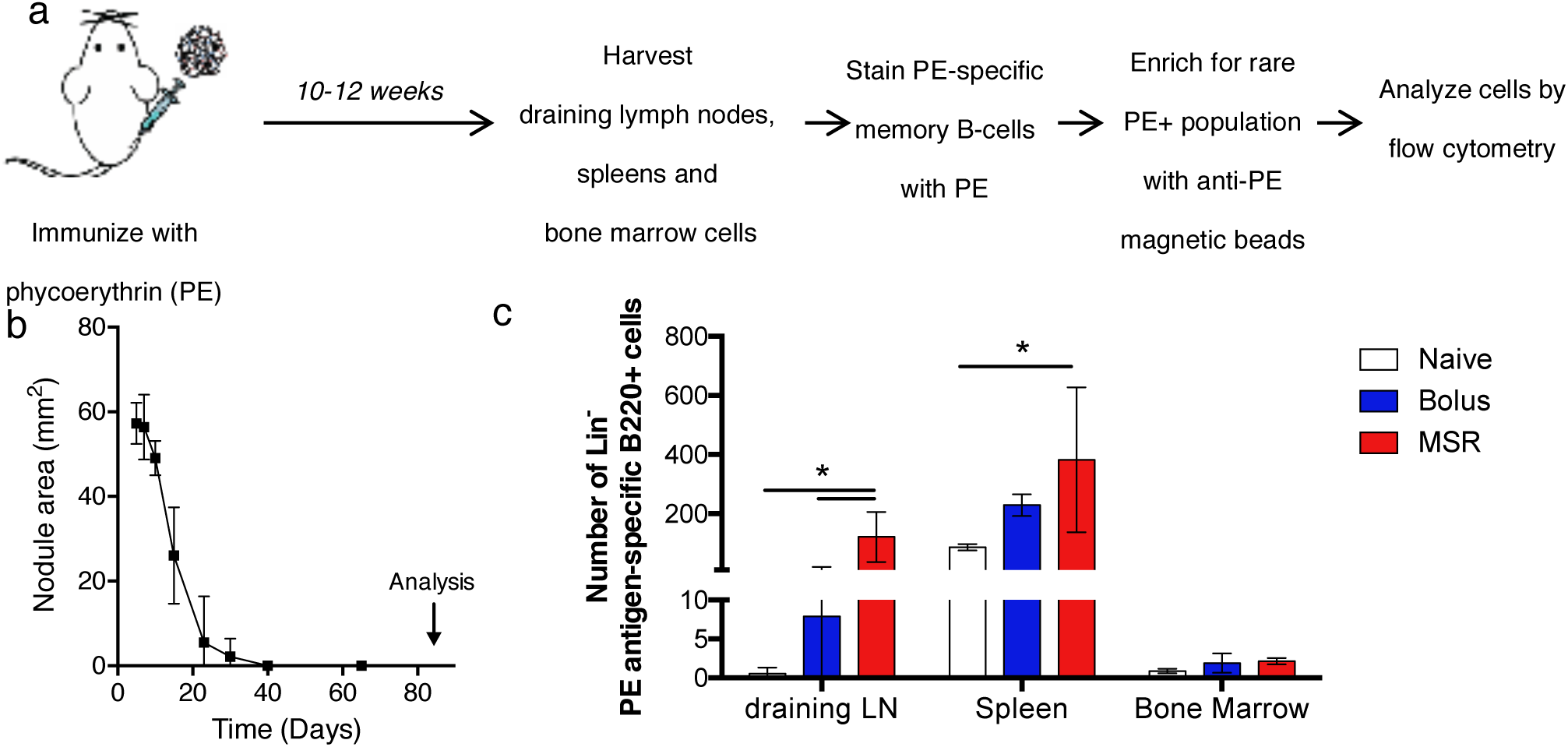
MSR vaccine improves memory B-cell generation. (a) Scheme of experimental design. Mice were left untreated (naïve) or received a phycoerythrin (PE)-containing vaccine in soluble (Bolus) or scaffold form (MSR). 10-12 weeks after immunization, when the vaccine components have cleared, cells of the draining lymph nodes, spleens and bone marrow were stained with PE to tag PE-binding cells. Anti-PE magnetic beads were used to enrich for the are PE-specific memory B-cells and further stained for analysis by flow cytometry. (b) Scaffold size over time, data represents mean and SD, n=4. (c) Total number of PE-specific Lin-B220^+^ cells (Lin- represents CD4-CD8-F4/80-Gr1-Cd11c- live cells) in the draining lymph nodes, spleens and bone marrow of naïve and bolus or MSR vaccinated mice, 83 days after immunization. (data represents individual data point with mean), n=4. * indicates p<0.05.

### 2.4 MSR scaffold persistence and cell-recruitment is required for robust humoral response generation

To begin to investigate how the dynamics of immune cell recruitment and antigen availability impacted on the humoral response generated by MSR scaffolds, we determined the kinetics of antigen presentation and cellular infiltration in GnRH-OVA MSR vaccines. Following subcutaneous injection, the vaccine formed a dynamic nodule which peaked in size on day 7 and slowly degraded over the next several weeks. Scaffolds were undetectable 45 days after injection (Figure 5a). MSR scaffolds released antigen slowly with 27 ± 3% of the GnRH-OVA cargo desorbing in the first 12 hours followed by minimal release (10 ± 1% of total load) over 25 days *in vitro* (Figure 5b). To track antigen in scaffolds *in vivo*, we immunized mice with MSR vaccines formulated with fluorescent-OVA as the antigen (MSR-OVA*) and imaged the vaccine site 7, 14 and 30 days after injection using confocal microscopy. OVA* fluorescence was detected at all timepoints, with scaffold area decreasing after day 7, confirming scaffold dynamics and the persistence of antigen for up to 30 days (Figure 5c).

**Figure 5:**
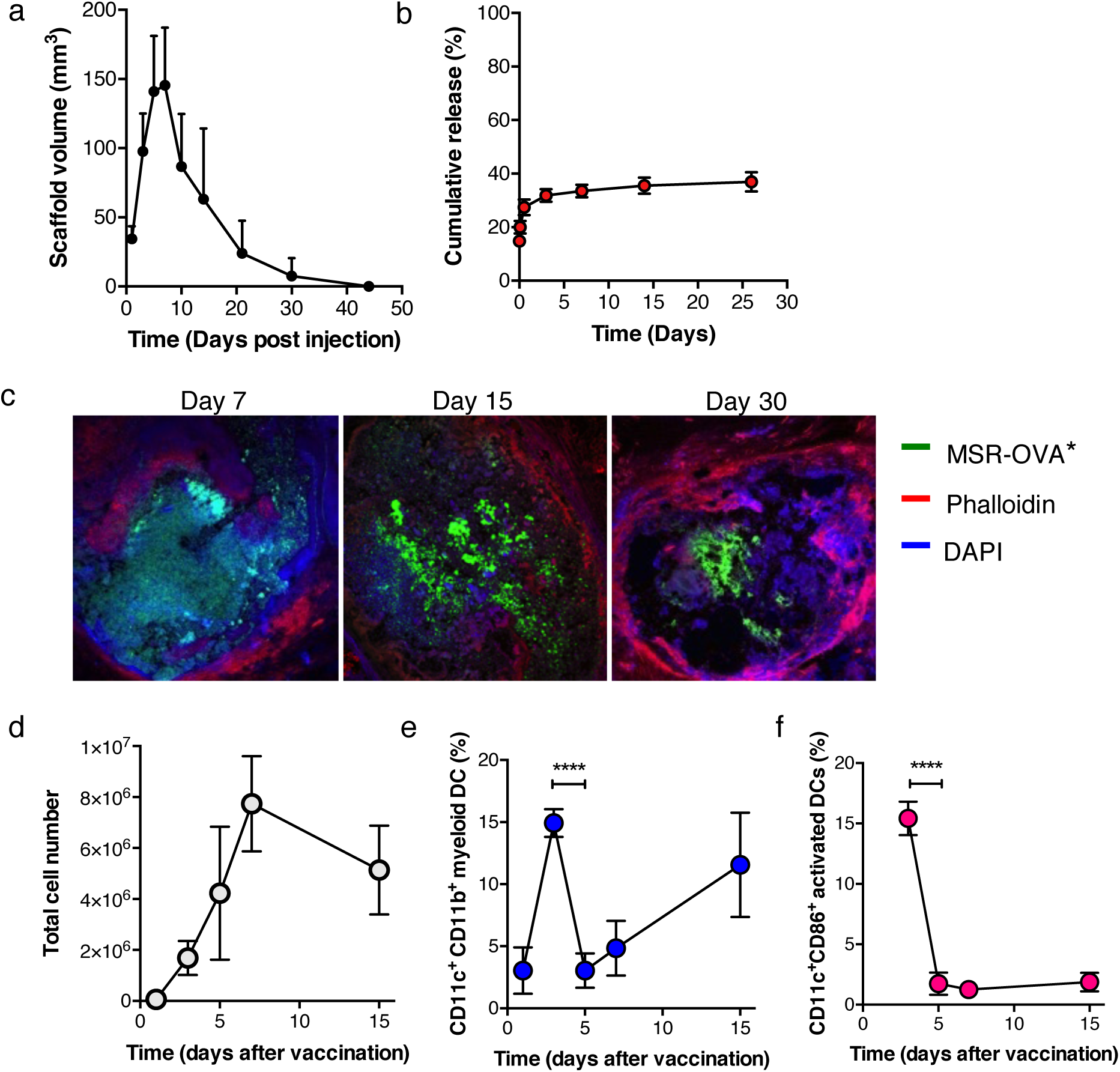
MSRs form dynamic scaffolds for sustained antigen release and DC recruitment and activation. (a) MSR vaccine scaffold volume over time. (mean and SD, n=4), (b) In vitro cumulative release of GnRH-OVA antigen from MSR scaffold. (mean and SD, n=3) (c) Representative confocal images of explanted MSR vaccine scaffolds containing fluorescently labeled OVA (OVA*) 7, 15 and 30 days after subcutaneous injection. Scale bar: 500µm. (d) Total number of cells in the MSR vaccine scaffold over time. (e) Percentage of CD11c^+^CD11b^+^ myeloid DCs in the MSR vaccine scaffold over time. (f) Percentage of CD11c^+^CD86^+^ activated DCs in the MSR vaccine scaffold over time (mean and SD, n=4), **** indicates p <0.0001

Next, the kinetics of cell recruitment by the MSR GnRH-OVA vaccines was quantified. The MSR vaccine has been engineered to recruit and activate host immune cells by incorporating and releasing GM-CSF and CpG-ODN ^[28]^. Consistent with our previous work, host immune cells were shown to infiltrate the MSR scaffold in a time dependent manner, peaking at day 7 (Figure 5d). CD11c^+^CD11b^+^ DCs were detectable at day 1 post immunization, their percentage peaked at day 3 and dropped sharply at day 5, followed by a secondary increase from day 7 to day 14 (Figure 5e). Recruited DCs were activated by the released CpG-ODN, as indicated by a higher percentage of CD11c^+^CD86^+^ DCs on day 3 after immunization (Figure 5f). The percentage of activated DCs dropped from day 3 to day 5, likely due to their homing to the draining lymph node (dLN). The percentage of activated DCs in the scaffold remained low from day 5 to day 14. Together, these data suggest the MSR GnRH-OVA vaccine induces rapid DC infiltration, activation and exfiltration, and serves as a site of sustained antigen presentation that may program immune cells for extended periods.

To investigate the relationship between the infiltrated immune cells and the anti-GnRH titer, the MSR vaccines were explanted at days 1, 3, 5, 7 and 15 after immunization (Figure 6a). Vaccination durations of less than 5 days resulted in no significant overall anti-GnRH IgG1 (Figure 6b) or IgG2a (Figure 6c) titers. Vaccines that were explanted on day 5 led to detectable amounts of IgG1 titer on day 14 and 20, but the response was transient and no titers were detected from this condition after 20 days. In contrast, a vaccination duration of 7 days or greater led to prolonged anti-GnRH IgG1 (Figure 6d) and IgG2a (Figure 6e) titers. Vaccines that were explanted on day 7 resulted in a significantly slower onset of titer production compared to vaccines that were explanted on day 15 and those that were not explanted. There was no difference in titer levels between vaccines that were explanted on day 15 and those that were not explanted. Together, this data suggest that vaccine duration of 15 days is necessary to generate both rapid onset and durable high anti-GnRH titers.

**Figure 6:**
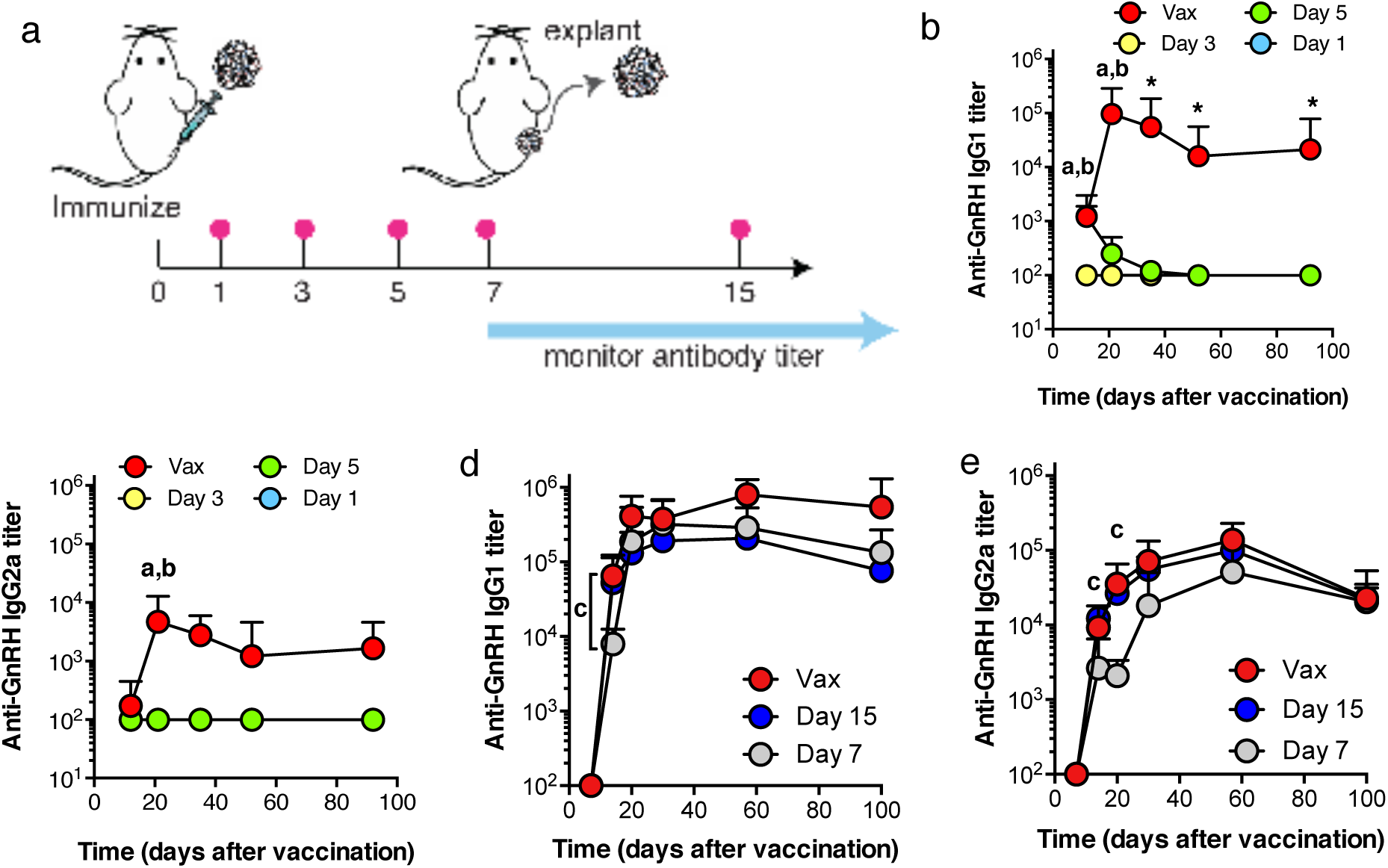
MSR vaccine duration impacts anti-GnRH antibody production. (a) Schematics of MSR vaccine regimen. Vaccine duration is varied from 1, 3, 5, 7 and 15 days, or indefinitely. (b-c) ELISA analysis of sera GnRH-specific IgG1 (b) or IgG2a (c) antibody from mice immunized with the MSR vaccine for a duration of 1 (Day 1), 3 (Day 3), 5 days (Day 5) or indefinitely (Vax). (d-e). ELISA analysis of sera GnRH-specific IgG1 (d) or IgG2a (e) antibody from mice immunized with the MSR vaccine for a duration of 7 days (Day 7), 15 days (Day 15) or indefinitely (Vax). (mean and s.d., n=4) a indicates p < 0.05 between Vax and Day 1, b between Vax and Day 3, c between Vax and Day 7 and * between Vax and all other groups.

To further probe the importance of cell recruitment for antibody generation, we created MSR vaccines with diminished cell recruitment capabilities. We previously reported that the compression of MSR scaffolds into a tablet (Monolith) decreased cell infiltration by reducing the 3D space available for immune cell infiltration ^[28]^. In addition, we encapsulated MSR solution into a nanoporous alginate hydrogel (MSR-Alginate) to prevent cell ingress (Figure 7a). When the modified GnRH-OVA MSR vaccines were used, they generated 1 to 2 orders of magnitude lower anti-GnRH titer than pristine MSR (Figure 7b).

**Figure 7:**
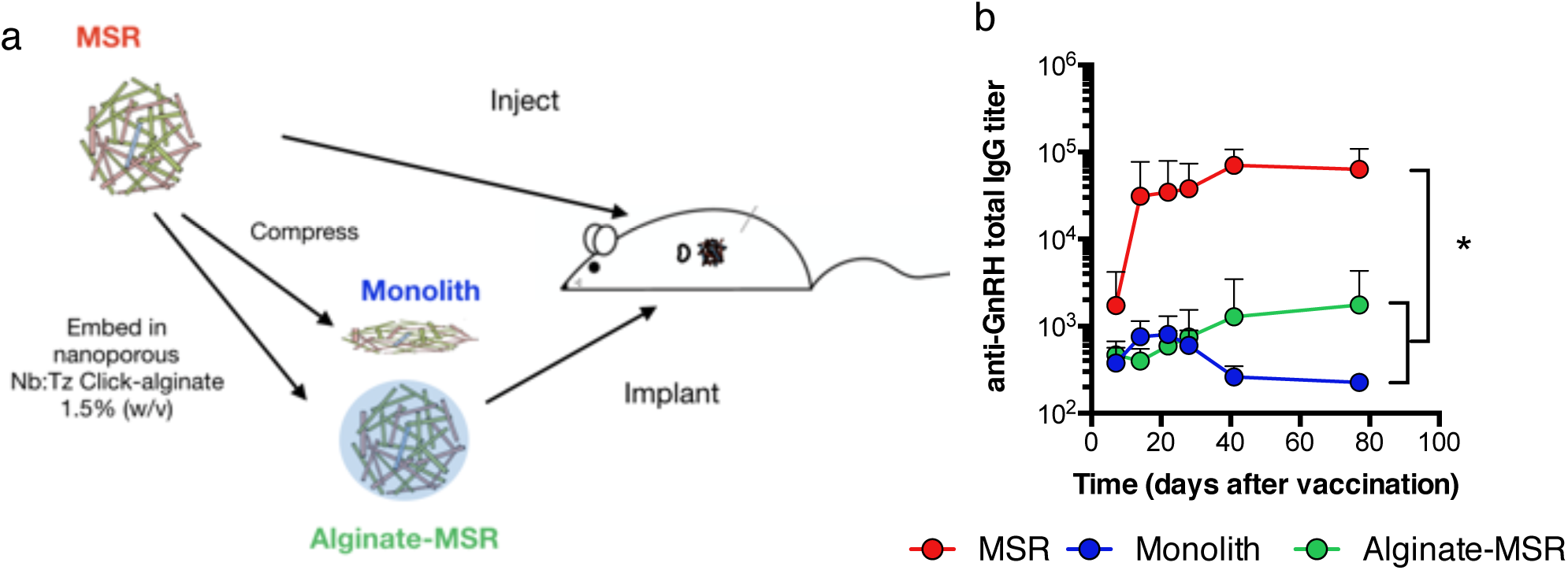
Reduced cell infiltration impairs antibody response. (a) Schematics of the modification of MSR vaccine to reduce cell infiltration. Lyophilized MSR vaccines were individually compressed into a 8mm-diameter disk using a lab press (monolith) or encapsulated into a nanoporous alginate hydrogel and implanted surgically in the subcutaneous space on mice’s flank (b) ELISA analysis of sera anti-GnRH total IgG titer over time. MSR condition indicates control MSR vaccine (not compressed nor incorporated into gel). All vaccine conditions contained the same masses of GM-CSF and antigen. (mean and SD, n=4), *, p<0.05 for all timepoints > 7 days.

### 2.5 Broad potential of MSR for vaccination against small molecules and linear peptide antigens

To demonstrate the broader potential of MSR vaccines, we tested their capacity to generate antibodies against different small molecule and linear peptide antigens (Figure 8a). Besides protein-derived antigens, there is broad interest in eliciting neutralizing antibodies against small molecules such as toxins or psychoactive substances to nullify their action ^[33,34]^. For example, vaccination has been investigated to produce antibodies that bind nicotine in the bloodstream to reduce the rewarding effect of tobacco and assist in smoking cessation ^[33,35]^. A single injection of MSR vaccines containing OVA-conjugated nicotine could induce high titers of anti-nicotine IgG, which remained elevated 6 months post injection (Figure 8b).

**Figure 8:**
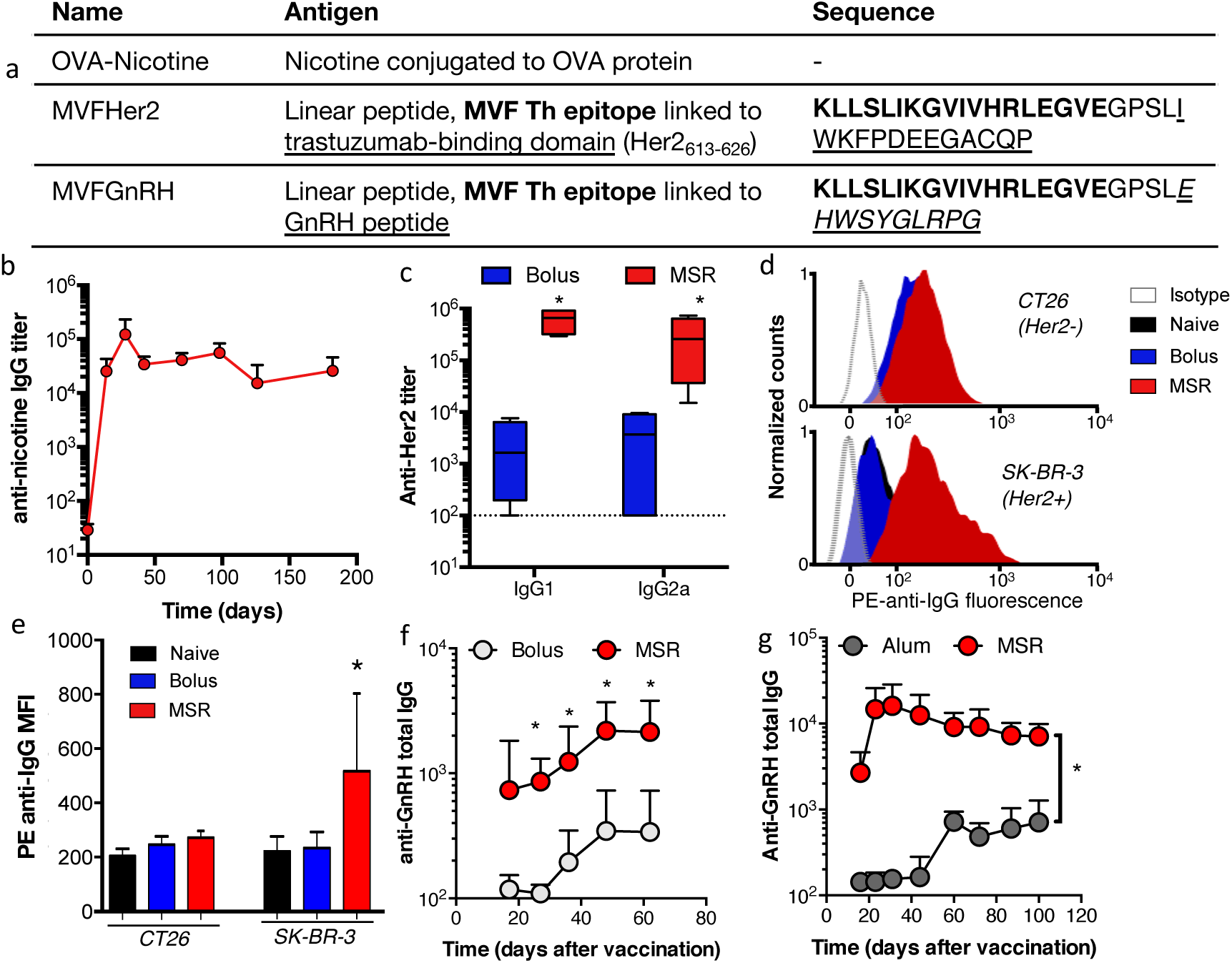
MSR scaffold vaccines induce robust humoral responses against a wide variety of small antigenic therapeutic targets. (a) Details of small molecule or linear peptide antigens used in vaccination studies. (b). ELISA analysis of anti-nicotine total IgG titer over time following vaccination using MSR vaccine. (c) ELISA analysis of anti-Her2 antibody on day 23 after vaccination with bolus (Bolus, n=4) or MSR vaccine (MSR, n=5); containing MVFHer2. Dotted line represents antibody levels in naïve mice; data represents mean and SD, * indicates p<0.05. (d-f) Flow cytometry analysis of serum IgG binding to non-Her2 expressing CT26 cells or Her2 overexpressing SK-BR-3 cells. (d) Representative anti-IgG fluorescence after treatment with serum from untreated (naïve) mice or animals immunized with a bolus (Bolus) or MSR (MSR) vaccine containing MVFHer2. (e) Quantification of IgG binding to CT26 and SK-BR-3 cells incubated with the serum of naïve (Naïve, n=5) or MVFHer2 vaccinated mice (Bolus, n=4 or MSR, n=5) * indicates p<0.05. (f-g) ELISA analysis of serum anti-GnRH IgG over time comparing mice immunized with a bolus (bolus, n=4) or MSR vaccine (MSR, n=4) containing MVFGnRH (f) or (g) MVFGnRH formulated in standard Alum adjuvant (Alum, n=4) and the MSR vaccine (MSR, n=4). data represents mean and SD; indicates p<0.05

Vaccines that generate antibody responses using carrier proteins may require cumbersome recombinant protein production and chemical conjugation processes, which can be costly, lead to batch-to-batch variability, and elicit off-target antibody responses against other T and B cell epitopes on the carrier protein itself ^[36]^. To circumvent this issue, we designed peptide constructs that incorporate a promiscuous T-helper cell epitope derived from measles virus fusion protein (MVF), a flexible linker, and the B-cell epitope of interest. In a first example, we targeted the oncogenic protein HER2/neu, a surface receptor that has been successfully targeted by monoclonal antibodies for the treatment of Her2^+^ breast and gastric cancer ^[37]^. To specifically induce therapeutic antibodies, we used Her2613-626 as the B-cell epitope, as it is within the portion of the protein bound by trastuzumab (Herceptin), a clinically approved monoclonal antibody that antagonizes the function of the Her2 receptor. Compared to a bolus, the MSR vaccine significantly enhanced both the IgG1 and IgG2a titer generated against the Her2 protein (Figure 8b). Moreover, serum IgG from MSR mice could bind to the Her2^+^ SK-BR-3 breast cancer cell line but not to CT26 Her2-cells. In contrast, serum from a bolus vaccine showed no detectable binding to Her2-expressing cancer cells (Figure 8d-e).

Finally, we investigated the anti-GnRH response generated against a synthetic MVFGnRH peptide antigen, comparing our MSR technology with a bolus vaccine and MVFGnRH formulated in the gold standard adjuvant Alum. In both cases, the total IgG titer induced by the MSR was on average an order of magnitude higher following a single injection (Figure 8f and g). Overall, this data illustrates the potential of MSR as a vaccine platform to elicit robust antibody responses against small antigens with otherwise low immunogenicity.

## 3. Discussion

The findings of this study demonstrate that a single injection of the MSR vaccine elicits potent and durable serum antibody titers against small antigens, including small molecules and short peptides. Both peptide-carrier protein constructs (e.g. GnRH-OVA and GnRH-KLH) and tandem peptide constructs (e.g. MVFGnRH and MVFHer2) demonstrated that the MSR vaccine induced more potent and durable humoral responses than traditional vaccinations. Notably, the MSR GnRH-OVA vaccine generated high anti-GnRH titers even 12 months after vaccination. Although carrier proteins can significantly increase immune responses against a peptide, this approach has several limitations. First, chemical coupling of specific residues on the carrier protein and the peptide can be difficult to control, and may lead to variability in vaccine response ^[38]^. Second, because the carrier protein contains multiple antigenic epitopes, there is likely skewed clonal dominance of carrier-specific B cell responses ^[36,39]^. For these reasons, tandem peptide constructs containing a single CD4 T-helper epitope and the target peptide were also explored. The MSR MVFGnRH vaccine produced 10-20 fold higher antibody titers compared to the bolus formulations, and MSR vaccination led to faster antibody production and overall higher response compared to a conventional Alum adjuvanted vaccine. Incorporating the antigen into the MSR scaffold likely resulted in multivalent display of the antigen on the MSR pores and surface, which may facilitate BCR crosslinking and enhance B cell activation for stronger humoral responses.

The MSR vaccine generated antibodies against a portion of the Her2 epitope that exhibited immunoreactivity to the native Her2 structure on a tumor cell surface. While Trastuzumab can inhibit tumor growth, mAb therapy lacks long-term efficacy due to the limited half-life time of the immunoglobulins, and repeated administration of the mAb results in many side effects ^[40–42]^. It has been demonstrated that Trastuzumab binds to the portion of the Her2 extracellular domain spanning between residues 563-626 that contains three loops ^[43]^. Previous studies identified a number of peptides that mimic portions of the Her2 domain at the Her2/Trastuzumab interface ^[44–46]^. Repeated vaccination using these peptides led to antibodies that could recognize Her2/neu displayed on a cell surface and induced Her2 internalization in a similar manner as Trastuzumab. Here we chose a short and linear peptide Her2_613-626_ to evaluate the robustness of the MSR vaccine. Other studies using the same peptide construct showed that multiple vaccinations were required to raise high titers ^[46]^. In comparison, a single injection of the MSR vaccine was able to induce antibody titers up to 2 orders of magnitude higher than the bolus vaccine. In particular, the antibodies recognized Her2 displayed on human breast tumor cells, whereas the antibodies produced by the bolus vaccine did not show any significant binding. Although linear peptides are highly flexible and inexpensive to synthesize, they can adopt a variety of conformations in buffer and only a subset of these conformations are ultimately responsible for antibody reactivity. Therefore, future studies should explore various cyclic peptides mimicking the Her2 loop to confer antibody specificity, and evaluate the anti-tumor efficacy. Overall, these data suggest that the MSR vaccine can likely promote continuous *in situ* antibody production, generate focused and robust humoral anti-tumor immunity and form long-term immunological memory.

The relationship between MSR vaccine duration and the kinetics and maintenance of the humoral response suggests the importance of prolonged DC programming in the MSR vaccine. The DC profile in the MSR vaccine showed two distinct waves; the first one peaked on day 3 and rapidly diminished by day 5, and the second initiated after day 7. The first wave of DCs was likely due to recruitment mediated by release of GM-CSF. The residence of these DCs in the vaccine was transient, likely due to activation by the released CpG-ODN and homing to the dLN. The second wave of DCs was possibly a result of cytokines and chemokines secreted by the first wave of DCs and other immune cells. Interestingly, the first wave of DCs exhibited an activated phenotype, while the DCs in the second wave were mostly immature. Explanting the MSR vaccine between day 1 and day 5, before the first wave of DCs could be accumulated and subsequently home to the dLNs, resulted in no significant humoral response. When the vaccine was explanted after day 7, the vaccine effectively generated and maintained potent humoral responses over 100 days. Studies have shown that prolonged durations of vaccination (> 16 days) using engineered biomaterial scaffold vaccines significantly augmented persistent CTL responses locally and slowed tumor progression ^[47]^. Here, we showed that vaccine duration was also important in humoral responses. Explanting the scaffolds not only impacts local APC activation by MSR but also suppresses antigen delivery to draining lymph nodes. However, when the capacity of the MSR scaffold for cell infiltration was reduced (using MSR monolith or alginate encapsulation) and antigen presence maintained, the antibody was also partially diminished. This data suggest that the high antibody titer generated by MSR are, in part, generated by APC recruitment and trafficking through the scaffolds.

The enhanced humoral responses generated by the MSR vaccine correlated strongly with persistent GC activity. After encountering an antigen, activated B cells form GCs where they switch their immunoglobulin constant region from IgM to IgG, IgA or IgE, and undergo somatic hypermutation in the variable region to produce antibodies with high specificity and avidity ^[32]^. A number of biomaterial-based vaccine systems have been shown to stimulate persistent GC activity. For instance, poly(lactic-co-glycolic acid) (PLGA) nanoparticles containing multiple TLR ligands induced active GCs for ∼45 days, and multilamellar lipid vesicle based LN-targeting nanoparticles showed superior GC formation compared to bolus formulations on day 14 ^[48,49]^. However, these strategies required multiple injections to achieve this effect. In contrast, only a single injection of the MSR vaccine induced persistent GC activity for over 30 days. Specialized B cells that emerge from the GC secrete high affinity antibodies and can also differentiate into long-lived plasma and memory cells, which are necessary to generate life-long antibody production and rapid immune re-activation upon antigen re-encounter, respectively^[31]^. Consistent with the sustained GC responses, MSR vaccines generated long-term antibody production and higher numbers of memory B-cells than their bolus counterparts. Overall, this data illustrates to the potential of the MSR vaccines to generate both therapeutic and prophylactic antibody responses. The duration of the GC activity after immunization with the MSR vaccine also correlated with the amount of time the MSR vaccine remained visible at the injection site. Future studies could explore how varying MSR vaccine degradation kinetics impacts GC activity and antibody production. In addition, the kinetics of antigen presentation after vaccination with the MSR system could likely be further optimized, as recent studies have demonstrated a dramatic impact of the antigen presentation profile on the immune response ^[50,51]^. The role of persistent antigen presentation by different subtypes of APCs is also an important topic for further study.

## 4. Conclusion

The findings of these studies suggest the MSR vaccine may have high utility as a single-injection platform to generate potent humoral responses. We demonstrated that MSR vaccines induce high and persistent antibody titers against small challenging antigens with relevance to a variety of therapeutic applications including addiction therapy, immune-castration for feral animal population control and cancer treatment. Different subunits antigens were and can easily be swapped using the MSR system, which could therefore have significant impact in other areas, such as chronic bacteria or viral infection, and neurodegenerative diseases.

Compared to conventional vaccination strategies, MSR scaffolds improved the magnitude of antibody response, boosted GC persistence, and increased memory B-cell generation, all desirable features of efficient immunization schemes. The magnitude of the humoral response generated was dependent on scaffold persistence and local cell recruitment, which could be further modulated to refine MSR vaccines. These findings have broader implications for our understanding of current vaccines and the design of new vaccine technologies.

## 5. Experimental Section

### Peptides and proteins

All peptides used in this study were synthesized at least 95% purity from Peptide 2.0. Peptide sequences are as follows: CGnRH (CEHWSYGLRPG), scrambled CGnRH peptide (CRSYGPLHEWG), MVFGnRH (KLLSLIKGVIVHRLEGVEGPSLEHWSYGLRPG), MVFHer2 (KLLSLIKGVIVHRLEGVEGPSLIWKFPDEEGACQPL). Endotoxin free ovalbumin was purchased from Invivogen (vac-pova), and maleimide activated KLH was purchased from ThermoFisher Scientific (77605).

### MSR synthesis and MSR vaccine formulation

MSRs were fabricated as described previously. Briefly, 4g of P123 surfactant (average Mn ∼5800, Sigma-Aldrich) were dissolved in 150 g of 1.6M HCl solution and stirred with 8.6 g of tetraethyl orthosilicate (TEOS, 98%, Sigma-Aldrich) at 40 °C for 20 h, followed by aging at 100 °C for 24 hours. TEOS was extracted in 1% EtOH in HCl at 80 °C for 18 hours. To prepare the MSR vaccines, 4mg of the MSR were adsorbed with 100µg murine class B CpG-ODN (sequence TCCATGACGTTCCTGACGTT, IDT) and varying amounts of antigen (peptide or peptide-carrier protein conjugate) for 8 hours at room temperature under shaking, and subsequently lyophilized. Separately, 1mg of the MSRs were loaded with murine GM-CSF (Peprotech) for 1 hour at 37 °C under shaking. The MSRs were combined and resuspended in cold PBS prior to immunization. To determine loading efficiency, MSR loaded with antigen were resuspended in PBS, centrifuged to pellet MSR and the supernatant was collected. Antigen concentration was determined by measuring the concentration of remaining antigen in supernatant using a BCA protein quantification kit (Pierce)

### Peptide protein conjugation

To conjugate CGnRH to ovalbumin (OVA), OVA was first reacted with 30 molar excess of sulfo-SMCC (ThermoFisher Scientific) in PBS for 1 hour at room temperature. Subsequently, the maleimide-OVA was desalted to remove the excess sulfo-SMCC. Separately, CGnRH or CHer2 was reduced using 2 molar excess of TCEP (ThermoFisher Scientific) for 1 hour at room temperature. 100µg of Maleimide-OVA was then reacted with 100µg the CGnRH peptide in PBS for 12 hours at room temperature under shaking.

### Nicotine protein conjugation

Ovalbumin (OVA) was reacted with succinic anhydride (Sigma) for 2h in 0.1M borate buffer at a 1:50 molar ratio (1:2 amine to succinic anhydride). The resulting product was buffer-exchanged to MES buffer, pH 5.8 and reacted with 600 molar excess of 1-Ethyl-3-(3-dimethylaminopropyl)carbodiimide (EDC, Sigma) and 200 molar excess racemic aminomethylnicotine for 20h. Finally, the OVA-Nicotine product was buffer-exchanged to PBS for use in vaccination studies. Bovine Serum Albumin (BSA)-Nicotine conjugates were produced using the same procedure to detect anti-nicotine serum antibody by ELISA.

### Immunization

Unless otherwise noted, all in vivo studies were carried out using female C57BL/6J mice (Jackson Laboratories) between 6 to 10 weeks old at the beginning of the experiment. MSR vaccines were resuspended in 150ul of cold PBS and injected, via an 18G needle, subcutaneously in the intrascapular region with the mouse under brief isoflurane anesthesia. We have demonstrated that the MSR vaccine can be injected using 23G needles, but we used 18G needles in these studies to be consistent with our previous studies. All animal studies were performed in accordance with NIH guidelines, under approval of Harvard University’s Institutional Animal Care and Use Committee.

### Sera titer analysis using ELISA

Peripheral blood was collected periodically after immunization. Sera samples were analyzed for IgG1 (BD Biosciences), IgG2a (BD Biosciences) or total IgG (Biolegend) levels using ELISA. Briefly, ELISA plates were coated overnight in 4 °C with 30ug/ml of GnRH, 10ug/ml Her2 or 5ng/mL BSA-Nicotine in PBS. ELISA was performed according to established procedures, and anti-GnRH or anti-Her2 titers were defined as the lowest serum dilution at which the ELISA OD reading was equal to OD value 0.2.

### Germinal center characterization

To analyze GC formation, dLNs were isolated on days 7, 14, 25 and 50 after immunization. Cells were enumerated, and stained with anti-mouse B220 (eBioscience), anti-mouse GL7 (eBioscience) and Rhodamine-PNA (Vector) for 15 minutes on ice. Cells were washed and assessed using flow cytometry (BD Fortessa or BD LSR-II).

### Lymph node histology

dLNs were fixed for 1 hour with 4% paraformaldehyde (Thermofisher Scientific), embedded in Tissue-tek OCT (VWR) and cryo-sectioned using a Leica CM1950 Cryostat. Various sections from one LN were stained with anti-B220 (eBioscience) and anti-GL7 (eBioscience) and visualized using confocal (Zeiss LSM 710).

### Cell isolation from MSR scaffolds explanted from animals

Scaffolds were excised on day 1, 3, 5, 7 or 15 after immunization. The tissues were processed through mechanical disruption and suspended in PBS. The resulting cell suspension was then filtered through a 40 μm cell strainer to isolate the cells from the larger sized MSRs. The cells and small remaining MSR particles were pelleted, washed with cold PBS, and counted (Beckman-Coulter). The portion of cells in the mixture of cells and small silica particles was accessed in SSC and FSC gating in flow cytometry (BD LSRII or BD Fortessa). Based on the counts from Coulter counter and the percentage of cells determined from FACS gating, the number of live cells in the MSR scaffolds could be calculated.

### Analysis of DC recruitment to MSR scaffolds

APC-conjugated CD11c (eBioscience), FITC-conjugated CD11b (eBioscience) stains were conducted for DC and leukocyte recruitment analysis, and APC-conjugated CD11c and PE-conjugated CD86 stains were conducted for DC maturation analysis. Cells were stained with 7-AAD and fluorophore conjugated antibody for 15 minutes on ice, washed thoroughly and analyzed using flow cytometry (BD LSRII or BD Fortessa). Cells were first gated according to the viability channel, live cells were then gated according to positive FITC, APC and PE using isotype controls, and the percentages of cells staining positive for each surface antigen were recorded.

### Analysis of memory B-cells

Mice were immunized as described above. In brief, each mouse received a subcutaneous injection of 1µg GM-CSF, 100µg CpG and 30µg PE (R-PE, Thermosfisher) in PBS (Bolus) or loaded on MSR (MSR) as described above. 10-12 weeks later, mice were euthanized and their draining lymph nodes, spleen and a lower limb harvested. Lymph nodes and spleens were processed in to single cells suspensions as described previously ^[52]^. Marrow was flushed off cleaned bones using a 21G needle and further processed into a single cell suspension. As described previously[^32^], cells were stained with 1µg PE, enriched for PE^+^ cells using anti-PE magnetic beads (Miltenyi Biotec). Cells were then enumerated and stained with Pacific Blue anti-mouse CD4, CD8, F4/80, Gr-1, Cd11c, FITC anti-mouse B220 (eBioscience) and Near-IR Fixable LIVE/DEAD dye (Thermofisher). Cells were washed and analyzed by flow cytometry (BD LSR-II) to detect CD4^-^CD8^-^F4/80^-^Gr1^-^CD11c^-^B220^+^PE^+^ memory B-cells.

### Mass spectrometry

GnRH-OVA conjugate was synthesized as described above and desalted. OVA and the conjugates were analyzed using Agilent 6460 Triple Quadrupole Mass Spectrometer equipped with Agilent 1290 uHPLC.

### Preparation of modified MSR vaccine for reduced cell recruitment

MSR vaccines were formulated as described above with the following modifications. 200µg OVA conjugated to 100µg GnRH peptide (OVA-GnRH) was loaded onto per 4mg MSR, flash frozen in liquid nitrogen and lyophilized. 1µg GM-CSF was loaded per 1mg MSR. MSR monolith were prepare by mixing the GnRH-OVA/MSR lyophilized particles and the MSR/GM-CSF solution and compressing them into an 8mm table using a lab press. Gel-embedded vaccines (Alginate-MSR) were made using norbornene-modified alginate (Nb:Alg) and tetrazine-modified alginate (Tz:Alg) that were prepared as described previously ^[53]^. 1.12mg Tz:Alg and Nb:Alg were dissolved separately in MES buffer to 3%(w/v) and MSR vaccines was resuspended to a total of 75µl. All components were mixed and allowed to gel for 2h. MSR monolith and Alginate-MSR were surgically implanted in the subcutaneous space of mice.

### Statistical analysis

All values in the present study were expressed as mean ± S.D. Statistical analysis was performed using GraphPad Prism and Microsoft Excel. For serum antibody analysis, significance was determined using unpaired Mann–Whitney tests. For all other analyses comparing multiple groups, one-way ANOVA tests were performed. For all statistical tests, p-values < 0.05 considered significant.

## Acknowledgements

This work was funded by the National Institute of Health (R01 CA223255 and U54 CA244726), the Wyss Institute for Biologically Inspired Engineering, the Found Animals Foundation (D1213-W13), and a National Science Foundation Graduate Research Fellowship to AL. The authors would like to thank Charles Vidoudez and the Harvard Bauer Core Facility for their help with mass spectrometry, and Dr. Thomas Conlon, Dr. Soumya Badrinath for their helpful discussions.

