## Supplementary figures and images for "Single-shot Mesoporous Silica Rods Scaffold for Induction of Humoral Responses Against Small Antigens"

### Supplementary Figures

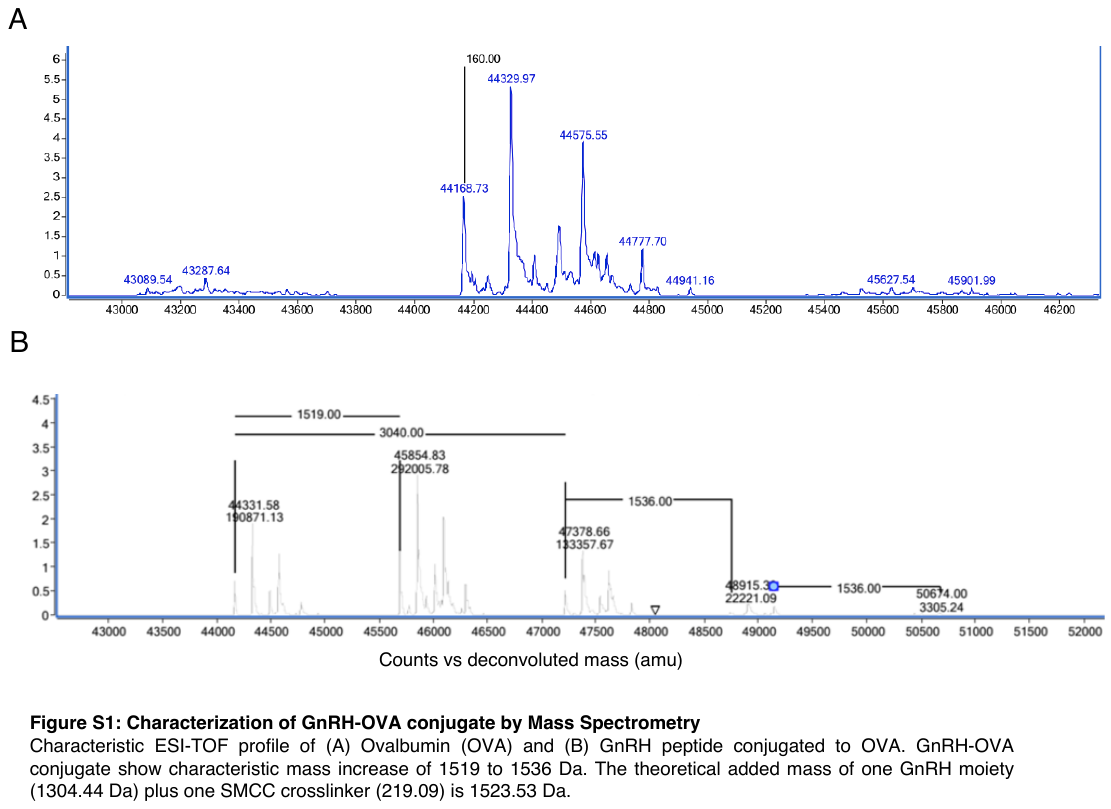

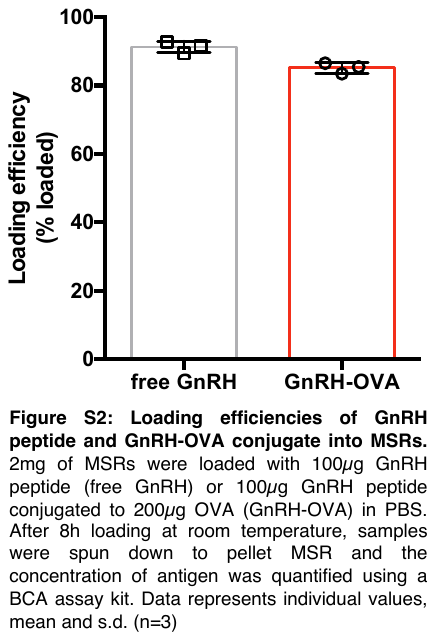

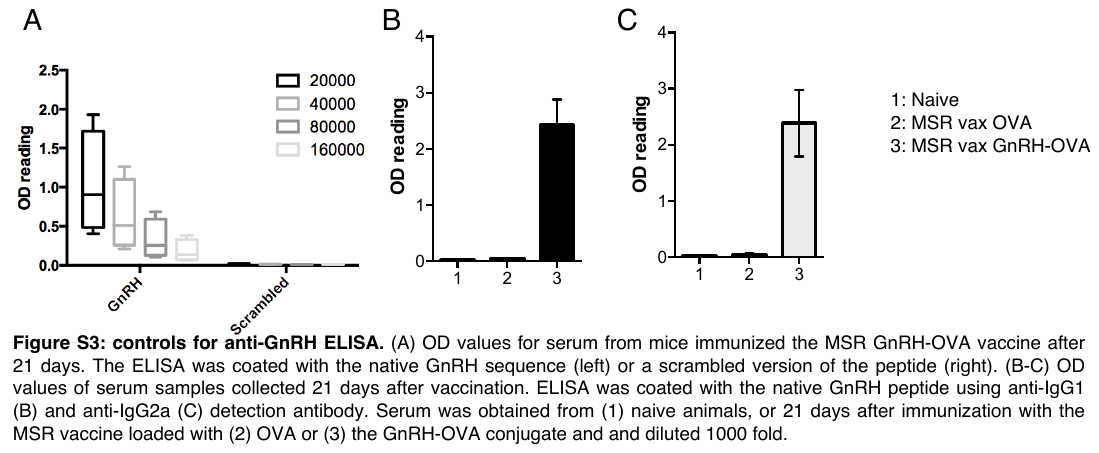

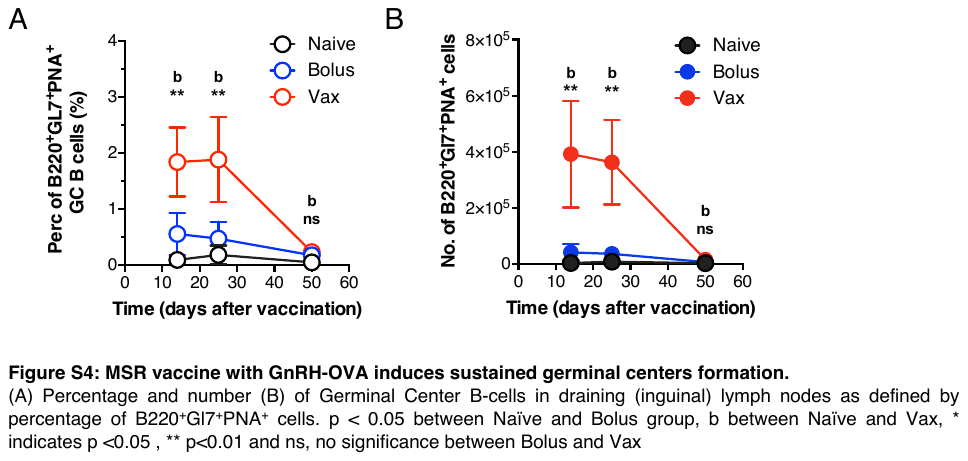

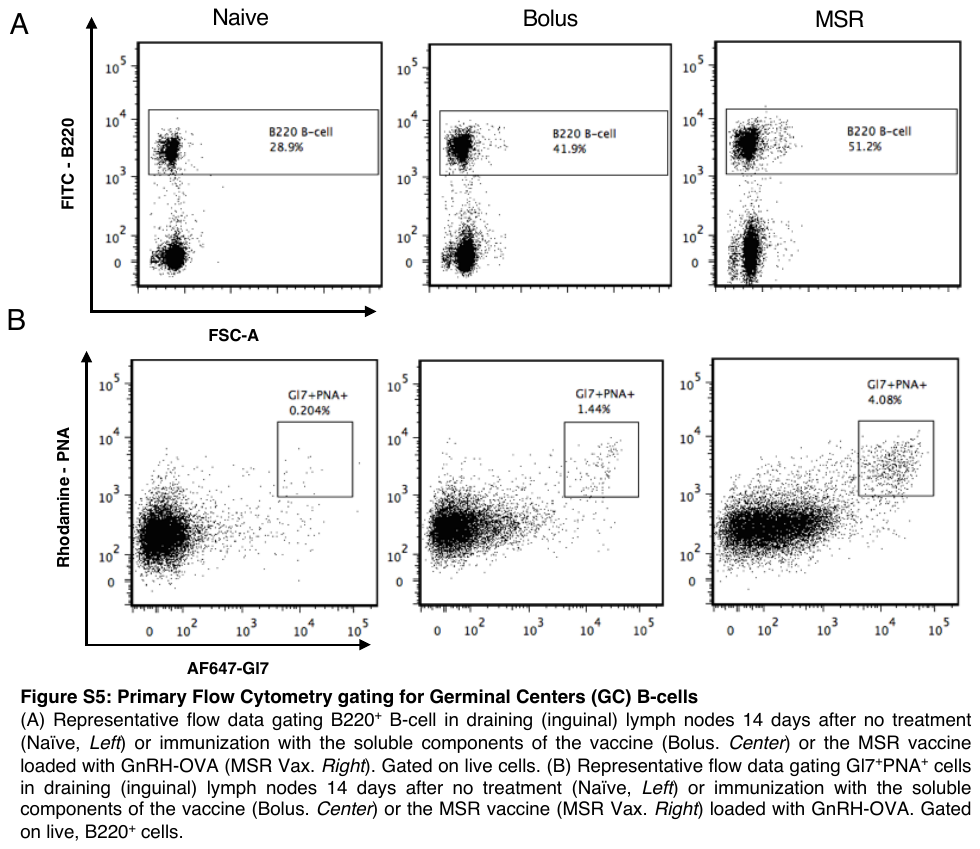

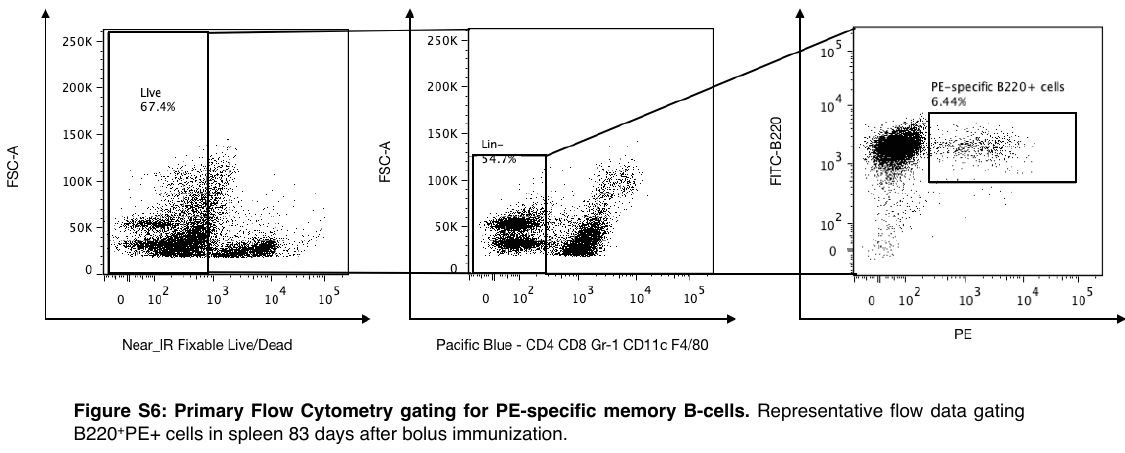
